# Estimation of age-specific susceptibility to influenza in the Netherlands and its relation to loss of CD8+ T-cell memory

**DOI:** 10.1101/259614

**Authors:** Christiaan H. van Dorp, Rutger G. Woolthuis, Jeffrey H. C. Yu, Rob J. de Boer, Michiel van Boven

## Abstract

The magnitude of influenza epidemics is largely determined by the number of susceptible individuals at the start of the influenza season. Susceptibility, in turn, is influenced by antigenic drift. The evolution of influenza’s B-cell epitopes has been charted thoroughly, and only recently evidence for T-cell driven evolution is accumulating. We investigate the relation between susceptibility to influenza, and antigenic drift at CD8^+^ T-cell epitopes over a 45-year timespan. We estimate age-specific susceptibility with data reported by general practitioners, using a disease-transmission model in a Bayesian framework. We find large variation in susceptibility, both between seasons and age classes. Although it is often assumed that antigenic drift drives the variation in susceptibility, we do not find evidence for a relation between drift and susceptibility in our data. This suggests that other factors determining the variation in susceptibility play a dominating role, or that complex influenza-infection histories obscure any direct effects.

**Preface to this bioRχiv pre-print:** We are currently in the process of making this manuscript ready for re-submission, and are resolving some issues brought forward by our referees. Most importantly, we aim to better incorporate the co-circulation of the various influenza A and B subtypes during the different seasons, both in the estimation of susceptibility and antigenic drift.

## Introduction

Since the year 1970, the Dutch research institute NIVEL has been using a network of general practitioners (GPs) to estimate the weekly incidence of influenza-like illness (ILI) in the Dutch population (Donker, 2016). This has resulted in one of the longest ILI-consultation time series available worldwide. The time series contains information about influenza epidemiology, and has enabled analysis of, among other things, the effect of humidity, school holidays (Te Beest *et al.*, 2013), waning immunity (Xia *et al.*, 2005), the impact of vaccination (McDonald *et al.*, 2016), and the validity of models of antigenic evolution (Ratmann *et al.*, 2012). For many of these studies it is presumed that the time series contains implicit information on the immune status of the population.

Immunity to influenza is of humoral (B cells) and cellular (T cells) nature, where the B cells and the antibodies they produce have received most attention. IgA antibodies against epitopes on hemagglutinin (HA) and neuraminidase (NA) provide neutralizing immunity, from which influenza is escaping by means of antigenic drift and shift, as visualized by antigenic cartography (Lapedes and Farber, 2001; Smith *et al.*, 2004; Bedford *et al.*, 2014; Neher *et al.*, 2016). Well-matching vaccines can be efficient in preventing infection (Shubin *et al.*, 2016), but require frequent updates, and vaccine mismatches are not unusual (Berlanda Scorza *et al.*, 2016; Teirlinck *et al.*, 2015). Cytotoxic T-lymphocyte (CTL) responses against peptidic epitopes of influenza’s internal proteins can provide long-term protection (Van De Sandt *et al.*, 2015) because these epitopes are highly conserved. Conserved T-cell epitopes are believed to be the cause of the relatively low illness severity in adults during the 2009 H1N1 pandemic (Sridhar *et al.*, 2013), and the 1957 H2N2 pandemic (Epstein, 2006), although in both cases antibodies against the more conserved stem of HA could also have played a role. Not surprisingly, these T-cell epitopes are considered to be good targets for a long-lasting, universal vaccine (Berlanda Scorza *et al.*, 2016; Sridhar, 2016).

By cleverly comparing substitution rates within and outside epitope regions in human and swine influenza A virus (IAV), Machkovech *et al.* (2015) recently demonstrated positive selection inside CTL epitopes of the nucleoprotein (NP). We also studied the evolution of IAV on the level of CTL epitopes, and found that from 1968 onwards, the number of epitopes in H3N2 has been decreasing (Woolthuis *et al.*, 2016). Both results support the idea that IAV is not only adapting to the human population by escaping from humoral immunity, but also—albeit more slowly—from cellular immunity.

On the clinical level, T-cell mediated immunity is receiving increasingly more attention. Using animal models and human cohort studies, it has become clear that cellular immunity can be responsible for reduced viremia, or even asymptomatic infection (Hayward *et al.*, 2015; La Gruta and Turner, 2014). T-cell memory does not provide neutralizing immunity. However, the role of T-cell memory in asymptomatic infection, which in turn reduces infectiousness, leads us to suspect that cellular immunity can have a profound effect on the epidemiology of IAV. Furthermore, since CTL responses provide long lasting protection (Van De Sandt *et al.*, 2015), the effects of CTL escape can act on a longer time-scale than antibody-antigenic drift.

In this paper, we study the effect of CTL and antibody (Ab) immune escapes of influenza on its epidemiology. We need two types of data for this study: the fraction of susceptible individuals, and the rate of Ab- and CTL-antigenic drift. In order to find the fraction of susceptible individuals, we fit a deterministic ordinary differential equation (ODE) model for transmission to ILI-consultation data. Since this model is quite complex, as the parameters to be estimated can be age- and season-specific, we employ a Bayesian framework, using Markov chain Monte Carlo (MCMC) for parameter estimation. For the other type of data, the antigenic drift, we use the average between-season Ab-antigenic drift as measured by Bedford *et al.* (2014). As a measure of CTL-antigenic drift, we use the disappearance of epitopes, or “epitope loss”, as defined recently (Woolthuis *et al.*, 2016). The measure “epitope loss” can roughly be seen as the number of epitopes that disappear from the virus from one year to the next.

Since we expect any relation between susceptibility and antigenic drift to be confounded by age, we stratify the data and model by age class. Incidentally, this allows us to normalize the susceptibilities of the older age classes with the susceptibility of the youngest children, who are likely to be immunologically naive (cf. Epstein, 2006). In this manner, we correct for external factors shaping epidemic size, that differ between seasons, but have the same effect on the age classes. Such effects include virulence and weather.

Contrary to expectations, we find no significant relation between either measure of antigenic drift, and susceptibility. We discuss explanations for the lack of this relation. This seems to contradict some of the findings presented by Bedford *et al.* (2014), who do find some evidence for a relation between antibody-antigenic drift, and the size of the epidemic in a period of two decades in the USA. We find that in the computation of antigenic drift, correct timing of epidemic seasons is essential, and ignoring this may easily lead to false conclusions. Furthermore, we argue that complex individual infection histories may impair our ability to use antigenic drift as a predictor for epidemic hazard.

## Results

We first discuss the details of the data and our model. Then we validate some of our estimates against independent data, and subsequently our estimates are interpreted. Finally, we compare susceptibility with CTL- and Ab-antigenic drift.

### Data and model selection

An overview of the (aggregated) ILI data and the fitted model is given in Figure 1. From the first season (1969/70) to the last (2013/14), the fraction of reported ILI is gradually decreasing. This holds for both the epidemic peaks in the winters, and for the lower off-season weeks. There is a variety of potential explanations for this trend (Dijkstra *et al.*, 2009). First, the circulation of ILI causing pathogens could be slowly decreasing. For the elderly, this could be partially due to changes in vaccination policy (McDonald *et al.*, 2016). Second, it could be that people have become healthier, or the virus less virulent (Fleming and Elliot, 2008). Finally, it could be that the reporting tendency has decreased over the last 45 years, because people less frequently visit their GP when experiencing mild symptoms. The latter explanation is most simple, seems biologically plausible, and is incorporated in the current model. The probability that a person with ILI consults their GP is denoted *q*, and we model the downward trend in reporting by allowing the probability *q* to depend on time (see Methods).

**Figure 1:**
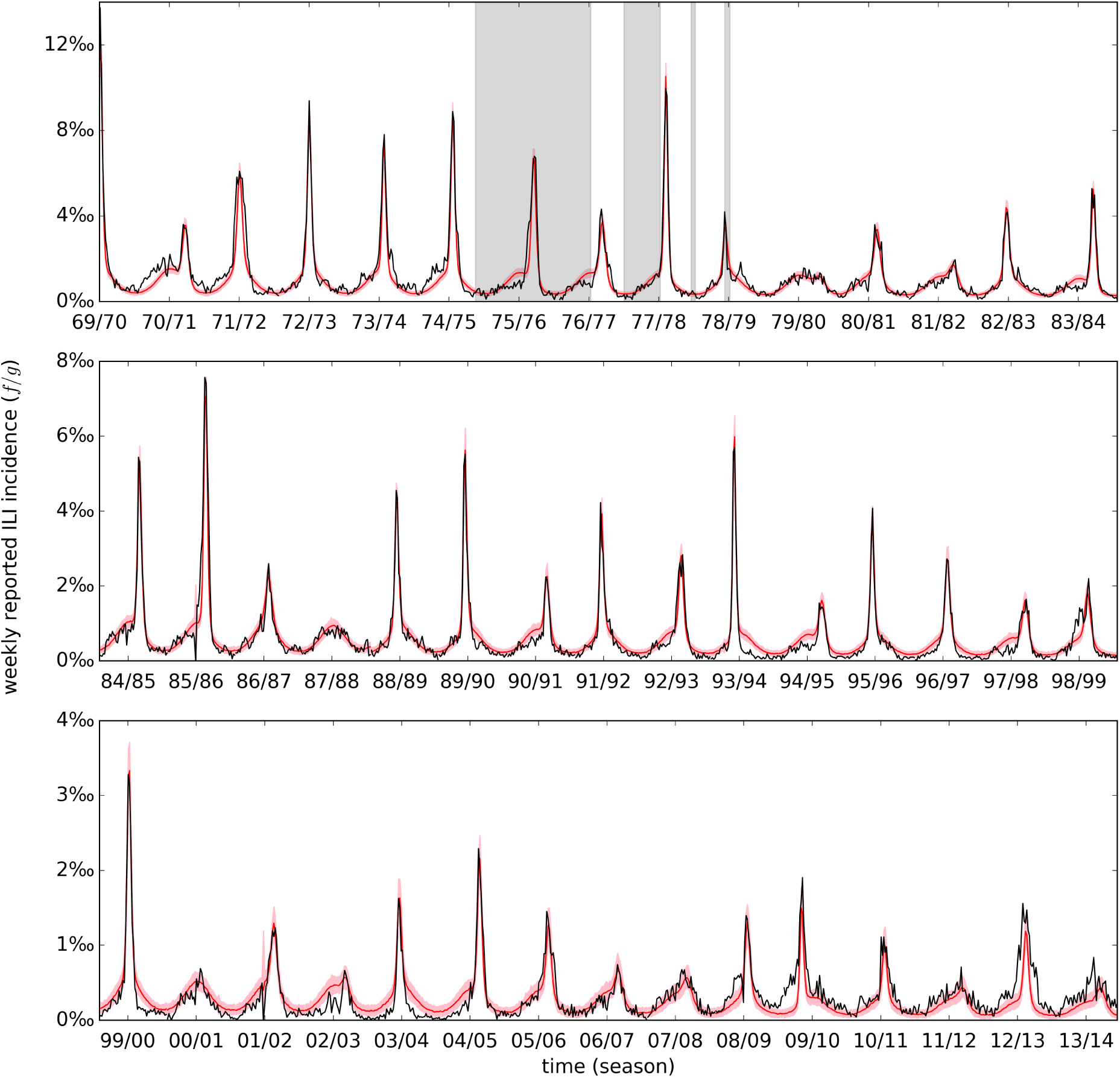
ILI data and model fit. Shown is the aggregated weekly ILI incidence (i.e., summed over the age classes), calculated by dividing the weekly number of reported cases (*f*) by the catchment population size (*g*) (black line), and the model fit (red line and pink band). The model fit is based on simulations using 200 samples from the posterior distribution of the baseline model. The red line represents the median incidence of the simulated data, and the pink band indicates the 2.5 to 97.5 percentile. The gray background color highlights weeks for which the age stratification and the catchment population size had to be imputed. Notice that the scale on the vertical axis is different for the three panels, as the reported incidence is declining.

An ILI data point consists of the number of reported ILI cases (*f*), and the catchment population size during that week (*g*, see Figure S1). The catchment population is a small sample from the entire Dutch population, and we assume that the total population is large enough so that the influenza epidemic can be modeled with a deterministic model of ODEs. ILI can be caused by a range of pathogens other than influenza (A and B), such as respiratory syncytial virus, rhinovirus, parainfluenza, and *Mycoplasma pneumoniae*. We refer to ILI that is *not* caused by influenza as background ILI.

The characteristic epidemic peaks are mostly caused by influenza, and are modeled with a transmission model (see below). Concerning the background ILI, a clear seasonal effect is visible (Figure 1), even during the ‘off-season’ weeks. For this reason, we use a descriptive model that takes seasonality into account. This approach is similar to the one taken by Van Noort **et al.** (2012). Furthermore, McDonald *et al.* (2016, 2017) used weekly laboratory surveillance data to discriminate between background ILI and influenza. The ILI in the ‘shoulder’ of the epidemics, turns out to be mainly caused by pathogens other than influenza.

Throughout, we stratify the **ILI** data into 6 age classes (Methods). The contacts between these groups are modeled using an age-specific contact structure that was derived with data from a human contact study (Mossong *et al.* (2008); see Data). The ODE model for influenza therefore consists of 4 x 6 compartments. For each age class, we have susceptible individuals (*S*), early and late stage infected individuals (*I*_1_, *I*_2_), and recovered individuals *(R;* not explicitly modeled). The two infection stages ensure a more realistic (Erlang) distribution of the length of the infectious period, and we assume no difference in infectiousness between the stages. The mean duration of the infection (1/γ) is not estimated; rather, we take 1/γ = 3.0 days, which is within the plausible range (Carrat *et al*, 2008).

For the initial conditions of the model, we need the fractions of susceptible individuals (So) at the beginning of the seasons. In our model the parameter So is the probability that an uninfected individual is infected upon contact with an infectious person, and becomes sufficiently ill to have a probability *q* to visit a GP. Individuals that do become infected, but experience mild disease (e.g. due to T-cell memory (Sridhar *et al.*, 2013; Epstein, 2006)), are therefore not considered to be susceptible, and are assumed to be hardly infectious, due to limited viral shedding (Laurie *et al.*, 2010). Notice that there is a slight mismatch between our interpretation of *S*_0_, and the usage of this parameter in the model. This discrepancy comes from the fact that it is hard to distinguish susceptibility from infectiousness in ODE models. For our purpose, both susceptibility and infectiousness are of interest. Hence, the current parameterization is suitable for our goal. This issue is discussed in more detail in the Supplementary material.

We use a mixed effects model for the susceptibility parameter *S*_0_. That is, *a priori*, the *S*_0_ are assumed to be sampled from Logit-Normal distributions, with unknown mean (*µ*_*susc*_) and standard deviation (*σ*_*susc*_). We consider versions of the model where *µ*_*susc*_ and *σ*_*susc*_ are dependent on the age class, and also a version where µ_susc_ is a linear function of the season *s* (see Estimation of the parameters).

Several models are compared with information criteria WAIC and WBIC (Table 1 and see Methods). Our analyses show that age-dependent susceptibility and periodic background ILI are essential elements of the model. The model that incorporates an age effect for *S*_*0*_ is favored over a model where every *S*_*0*_ has the same prior distribution. The model incorporating a time effect for *S*_*0*_ fits the data well, but has too many (effective) parameters.

**Table 1:** Model selection. Models are compared with WAIC and WBIC (see Methods). The number of parameters used in each model is denoted by *k.* Each one of the models is defined by relations between the parameters (see Methods), defined by: 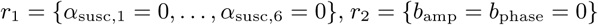, 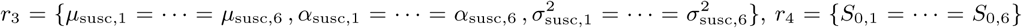. The best model (baseline) assumes relation *r*_1_, meaning that *S*_0_ is stationary. The baseline model has WAIC = 95 926 and WBIC = 97 073. For the model with constant background ILI we assume relation *r*_2_ in addition to r_1_, meaning that the background ILI is the same throughout the year. The model that only incorporates a single prior distribution for susceptibility (i.e. no age effect) assumes relation *r*_3_ in addition to *r_1_.* The model with age-independent susceptibility incorporates relation *r*_4_ in addition to *r*_1_, and (trivially) *r*_3_.

| scenario | relation | $k$ | $\Delta\text{WAIC}$ | $\Delta\text{WBIC}$ |
| --- | --- | --- | --- | --- |
| baseline | $r_1$ | 349 | - | - |
| trended susceptibility | - | 355 | 0.4 | 10.4 |
| single prior for susceptibility | $r_1, r_3$ | 339 | 8.9 | 304 |
| susceptibility independent of age | $r_1, r_3, r_4$ | 114 | 7285 | 5820 |
| constant background ILI | $r_1, r_2$ | 347 | 21 642 | 21 356 |

### Estimated reporting matches independent observations

In our analyses the reporting probability (*q*) is mainly a nuisance parameter that ties the observed number of ILI cases to the underlying circulation of influenza and background ILI. It is important, however, that estimates of the reporting do not systematically skew parameters that are truly of interest in this study. To provide external validation of our reporting rate estimation, we have used data from an independent study on influenza incidence (GIS, Friesema *et al.*, 2009, and see Data). Reassuringly, our estimates and those from GIS correspond well (Figure 2). Even the U-shape of the age-stratified reporting is captured by our estimates. Naturally, we can not use the results from this small number of consecutive years to extrapolate the validity of our reporting estimates to the entire time series (Figure S3), but the correspondence between the estimate of *q* and this independent data demonstrates our ability to extract this information from the ILI data.

**Figure 2:**
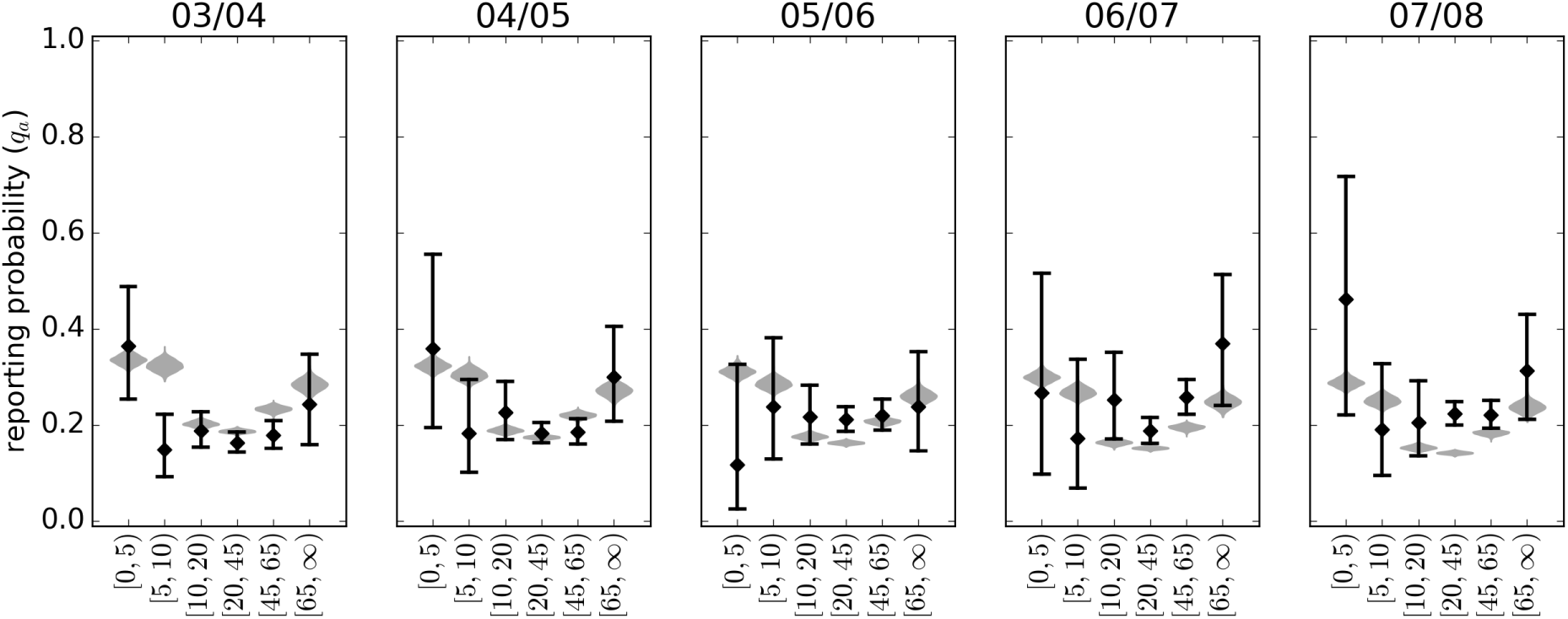
Reporting estimated from ILI data compares well to GIS reporting frequencies. The gray violin plots show the marginal posterior distribution of the reporting probabilities (*q*), which are used to scale reported ILI to influenza and background incidence. The black diamonds show the reporting frequency from the GIS data obtained from 2003 to 2008. The black whiskers indicate 95% confidence intervals (obtained using Jeffreys method).

### Susceptibility differs between and within age classes

The main outcome of the fitted Bayesian model is a set of estimates of the susceptibility to influenza of individuals in the 6 age classes at the beginning of 45 influenza seasons. Figure 3 shows an overview of these 6 x 45 estimates. Clear differences exist between the age classes (Figure 3B). The highest susceptibilities can be found among the elderly ([65, ∞) years), but this age class also shows the most variation and uncertainty. The elderly are followed by the youngest age class ([0,5) years) and the age class [45, 65) years. Children in the age class [5,10) years are least susceptible. On the other hand, age is not a perfect predictor for susceptibility, as the posterior predictive densities of the susceptibilities show a large overlap (Figure 3C). Summarizing, susceptibility is determined by both age and calendar time (epidemic season). The cause of the effect of age on susceptibility must be sought in individual health states and immunological history, and that of the epidemic season in antigenic drift, viral fitness, and environmental factors.

**Figure 3:**
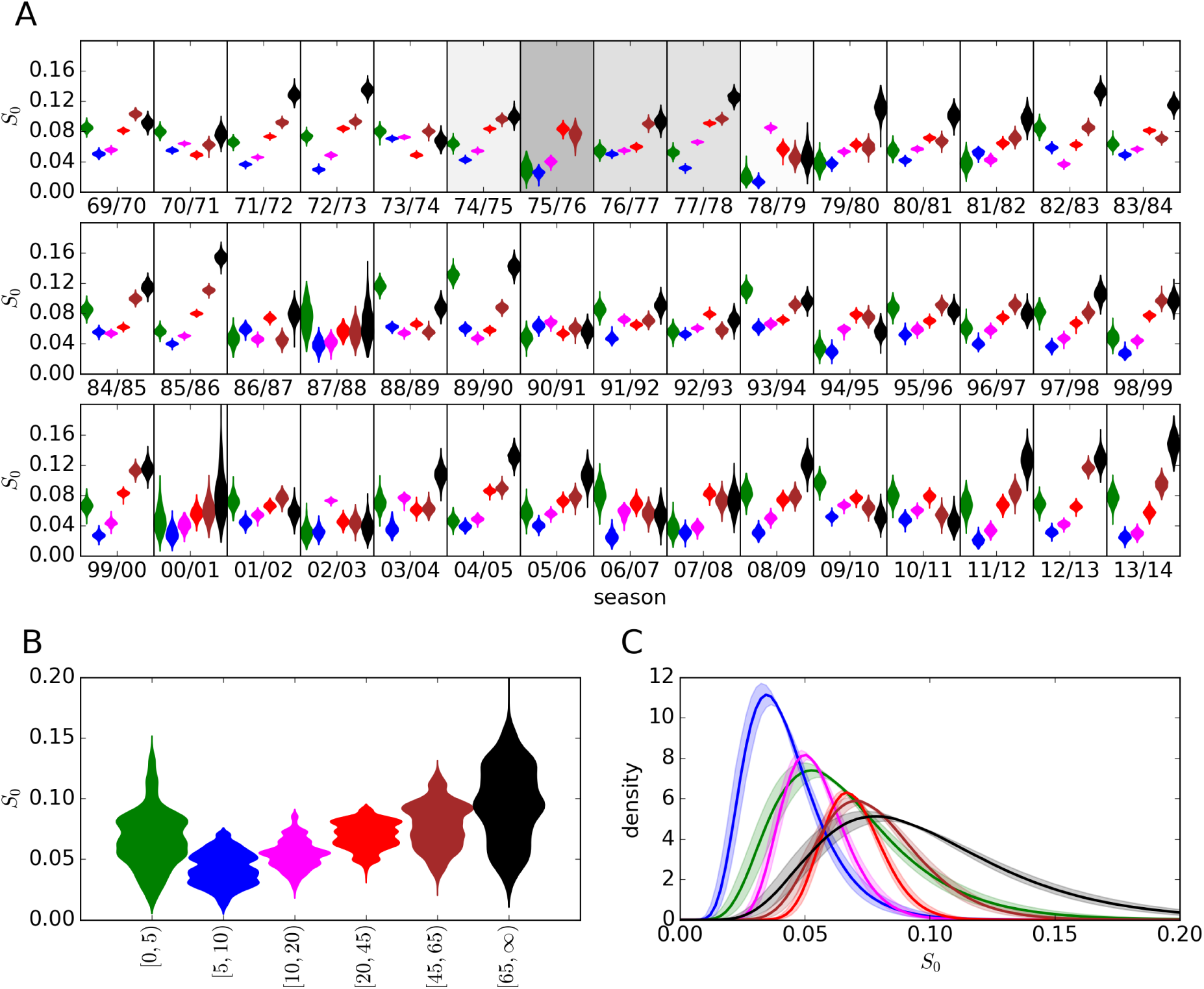
Susceptibility estimates stratified by season and age class. (A) The violin plots show the marginal distribution of the susceptibility parameters *S*_0,*a*_ for *a* = 1,…, 6. The color coding for age classes is as follows: green: [0, 5) years, blue: [5, 10) years, cyan: [10, 20) years, red: [20, 45) years, brown: [45, 65) years, black: [65, ∞) years. The gray background indicates missing age stratification during a season, with brightness proportional to the number of non-missing weeks (cf. Figure 1). (B) The samples from the posterior distribution are aggregated by age class. (C) Posterior predictive distributions of age-specific susceptibilities, i.e. the Logit-Normal distributions of the parameters *S*_0,_*_a_* are shown (*a* = 1,…, 6), with hyper parameters taken from the posterior distribution. The lines indicate the median densities, and the bands represent the interquartile range. The posterior modes and 95% CrIs of all susceptibilities are listed in Supplementary Table 1.

We observe considerable variation of susceptibility both within and between age classes. As we will argue now, this variation is not entirely random. By performing pair-wise comparisons between age classes over all seasons (Figure 4A), we make the following observations: (i) The susceptibility of closely related age classes is correlated significantly, or has a tendency to be correlated, except for the age classes [10, 20) and [20, 45) years. (ii) Susceptibility of children in the age class [0, 5) years is only somewhat correlated with age class [5,10) years. (iii) Susceptibility of adults and elderly is not, or negatively correlated with the susceptibility of the age classes [5,10) and [10, 20) years.

**Figure 4:**
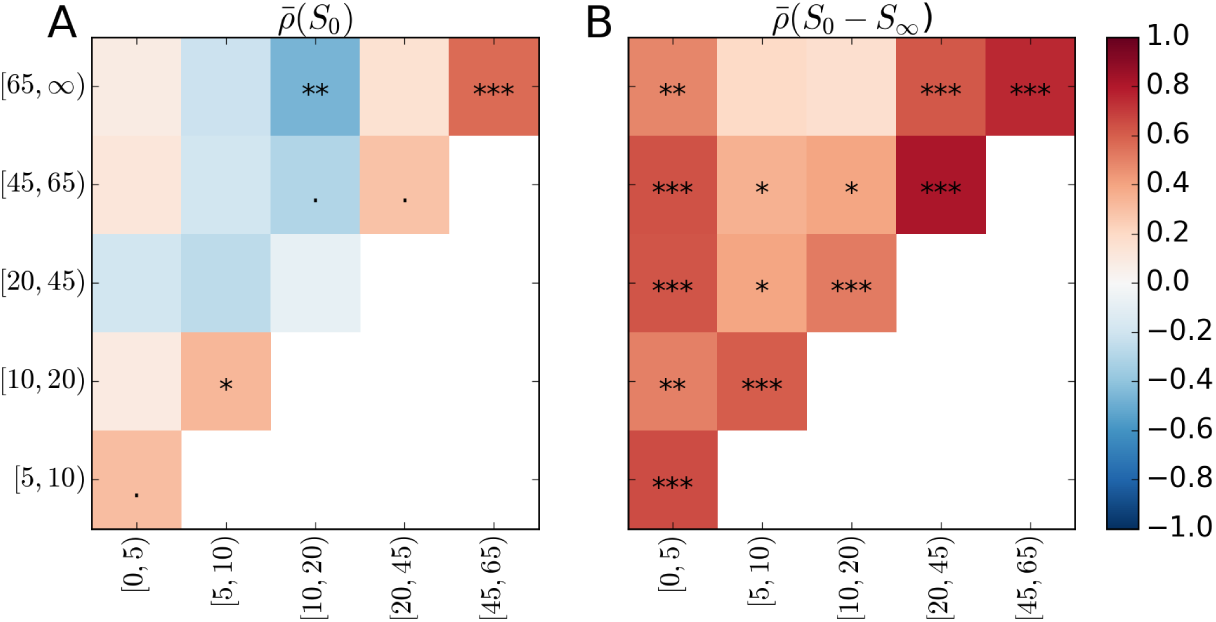
Correlations between estimated age-specific susceptibilities. (A) Every square is colored according to the expectation of the Spearman rank correlation between *S*_0 a1_ and *S*_0_ *_a2_* (with 1 *≤ a_1_ <* a_2_ *≤* 6). For instance, in seasons where individuals in the age class [45,65) are highly susceptible, the elderly (age class [65, ∞)) tend to be susceptible as well. The stars indicate the expectation of the significance level: ‘•’: 0.1 ≥ *p* > 0.05, ‘*’: 0.05 ≥ *p* > 0.01, ‘**’: 0.01 ≥ *p* > 0.001, ‘* * *’: 0.001 ≥ *p.* The seasons 1975/76 and 1978/79 are not included in the analysis, because of the missing age-stratification at the time of the epidemic peak. (B) Similar to (A), but then comparing the cumulative incidences of the age classes, i.e. the fraction of individuals that has been infected during the season (*S*_0_ – *S∞*, where *S∞* denotes the fraction remaining susceptible at the end of the epidemic).

These observations are understandable in light of immunological history. Individuals in closely related age classes are more likely to share a common infection history than individuals with a greater age difference, which would explain observation (i). Since children in the age class [0, 5) years are more likely to be immunologically naive, their susceptibility does not resemble that of older individuals, although a certain overlap is again to be expected with the age class [5,10) years, hence observation (ii). Observation (iii) is more difficult to explain, and might be related to “original antigenic sin” (OAS), or the related “antigenic seniority” (Lessler *et al.*, 2012). Recently, it was shown that antigenic seniority is likely to generate negative correlations between young and old birth cohorts with respect to susceptibility to severe avian influenza infection (Gostic *et al*, 2016). Such a mechanism could work more generally. If an influenza antigen resembles another recent antigen, then individuals in the age class [10, 20) years mount an effective immune response, and their susceptibility is low. Although older individuals also encountered the recent antigen, they may have failed to create memory against it (due to OAS), and suffer the consequence: increased susceptibility for currently circulating viruses. Vice versa, when older individuals respond well, this may be due to similarity to an old antigen (cf. the antigenic thrift model (Wikramaratna *et al.*, 2013)), which individuals in the age class [10, 20) years never encountered. Again, the fact that we find intuitive correlations between the estimated susceptibilities of the various age classes illustrates that we are obtaining meaningful estimates of *So* from the MCMC procedure.

An alternative measure for the immune status of the population is the attack rate (*S*_0_ – *S*_∞_, see Figure 4B). Arguably, it is more straight-forward to extract attack rates from ILI data than susceptibility. In fact, the attack rate (or average incidence) has been used in similar studies before (Bedford *et al.*, 2014). In the standard SIR model, the relation between susceptibility and attack rate is quite simple, albeit non-linear. The non-linearity can be described as herd immunity. Herd immunity becomes more important when multiple compartments (in our case age classes) are involved; the attack rate within a class becomes a poorer ‘predictor’ for the susceptibility of this class, since the other classes interfere. This can be seen in Figure 4B. In a way, estimating susceptibility with a compartmental model could be seen as a method to de-correlate age-stratified incidence data.

Using the estimates of *S*_0_ and the contact matrix, we can compute for each season the effective reproduction number *ℛ*_eff_ at the start of the epidemic (Figure 5), still assuming that individuals that experience mild disease hardly contribute to the epidemic. The effective reproduction number is related to the basic reproduction number in the sense that they are equal when there is no pre-existing immunity. On average, *ℛ*_eff_ equals 1.31 (95% CrI: [1.07, 1.49]; aggregated over all seasons). It is to be expected that the epidemic in a season with a high *ℛ*_eff_ starts earlier than epidemics in seasons with low *ℛ*_eff_, but we do not find any evidence for this in the data. We do find a strong correlation between the *ℛ*_eff_ of a season, and the precision with which we can estimate the start of the epidemic (Figure 5), as the interference of background ILI is stronger for small epidemics. Perhaps our inability to find a relation between timing and reproduction number is due to the lack of precision with which we can estimate this timing.

**Figure 5:**
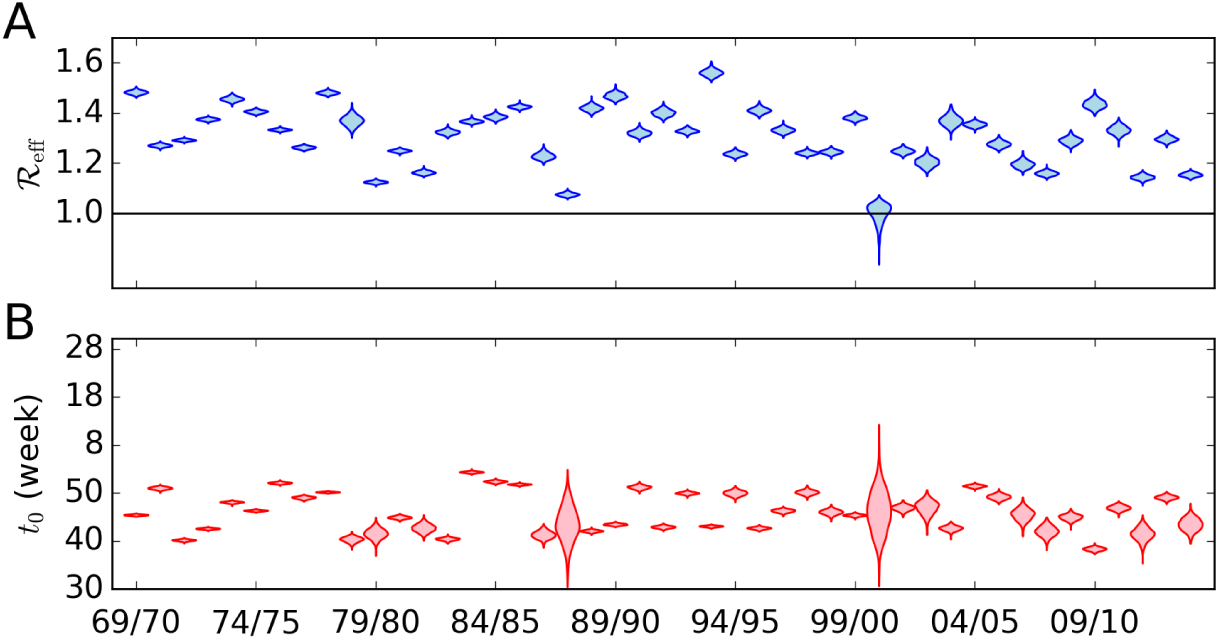
Estimates of the reproduction number and the onset of the epidemics. (A)The posterior densities of the compound parameter *ℛ*_eff_ (the effective reproduction number) for the 45 seasons. The horizontal black line indicates the epidemic threshold *ℛ*_eff_ = 1. (B) Estimates of t0, the start of the 45 epidemics (week number). The precision (sd(t0)^-1^) of the start of the epidemic is strongly correlated with the posterior mean of *ℛ*_eff_ (Spearman *ρ* = 0.85, *p* < 10^-12^). The posterior modes and 95% CrIs of *ℛ*_eff_ and t0 are listed in Supplementary Table 1.

Some of the estimated reproduction numbers can easily be related to the circulation of specific strains in the Netherlands. For instance, the relatively high reproduction number at the start of the 1993/94 epidemic season can probably be attributed to a strain from the BE92 (H3N2) cluster, which in antigenic space is far away from the previous BE89 cluster (Smith *et al.*, 2004). In addition, the cluster transition around the year 1993 coincides with a large CTL-antigenic jump (Woolthuis *et al.*, 2016). Similarly, the high *ℛ*_eff_ in 2009/10 is due to the pandemic H1N1 strain. The reappearance of H1N1 in 1977 (in the form of the A/USSR/90/77 strain) might be responsible for the high *ℛ*_eff_ in 1977/78. For 1969/70, we only have data for the second half of the epidemic. The fact that we are able to estimate susceptibility and a reproduction number for this season, is due to the mixed effects model for *S_0_.* The high *ℛ*_eff_ for this season corresponds well with estimates from England and Wales (Fleming and Elliot, 2008), and might be due to the re-assortment event that led to the replacement of H2N2 by H3N2. No H3N2 strains circulated during the 2000/01 season (Dijkstra *et al*., 2009), resulting in an estimated *ℛ*_eff_ around 1.

### CTL- and Ab-antigenic drift do not predict (relative) susceptibility

Previously, we calculated how many CTL epitopes appear or disappear from IAV between consecutive seasons (Woolthuis *et al.*, 2016, and see Data). Since the escape of an epitope should lead to loss of CTL memory in some individuals, we test whether the number of lost epitopes in H3N2 can explain susceptibility as estimated above. We focus on the H3N2 subtype, since this subtype circulates the most, causes the most severe disease, and evolves most rapidly. It is expected that an effect of CTL epitope loss on estimated susceptibility is small in the youngest children, because most of them have not been infected before, and tend to lack CTL memory against influenza. Additionally, the fact that CTL-epitope regions in IAV are relatively conserved (Machkovech *et al*., 2015), could be an indication of a fitness cost associated with mutations in these regions (Berkhoff *et al*., 2006). A strain that has many mutations in these epitopes, could therefore suffer from diminished infectiousness, when compared to the wild-type form in immunologically naive populations.

Apart from intrinsic fitness effects, variation of the susceptibility of the youngest children should also reflect environmental influences on influenza transmission, such as (indoor) humidity and temperature (Lowen *et al.*, 2007; Lowen and Steel, 2014). We normalize for such yearly effects by considering the susceptibility of the age classes, relative to the susceptibility of the youngest age class (henceforth “relative susceptibility”). In the following analyses, we leave out seasons dominated by H1N1 (1983/84, 1986/87, 2000/01, 2007/08, 2009/10, 2010/11). Likewise, seasons with missing age-stratified data around the epidemic peak we also removed from the analyses (1975/76, 1978/79).

We use the antigenic distance between strains (Bedford *et al.*, 2014) as a measure of Ab-antigenic drift. In short, this antigenic distance is derived from hemagglutinin inhibition (HI) assays using Bayesian multidimensional scaling (BMDS). The location of the strains along the first antigenic dimension is informed by a “drift prior”, in order to solve identifiability issues. Displacement along this axis is therefore a measure of antigenic drift. The antigenic drift between epidemic seasons is derived by first averaging the first coordinates of all strains in a particular year, and then taking the difference of the averages. No correlation between Ab-antigenic drift and relative susceptibility can be found (Figure 6A and Table 2).

**Figure 6:**
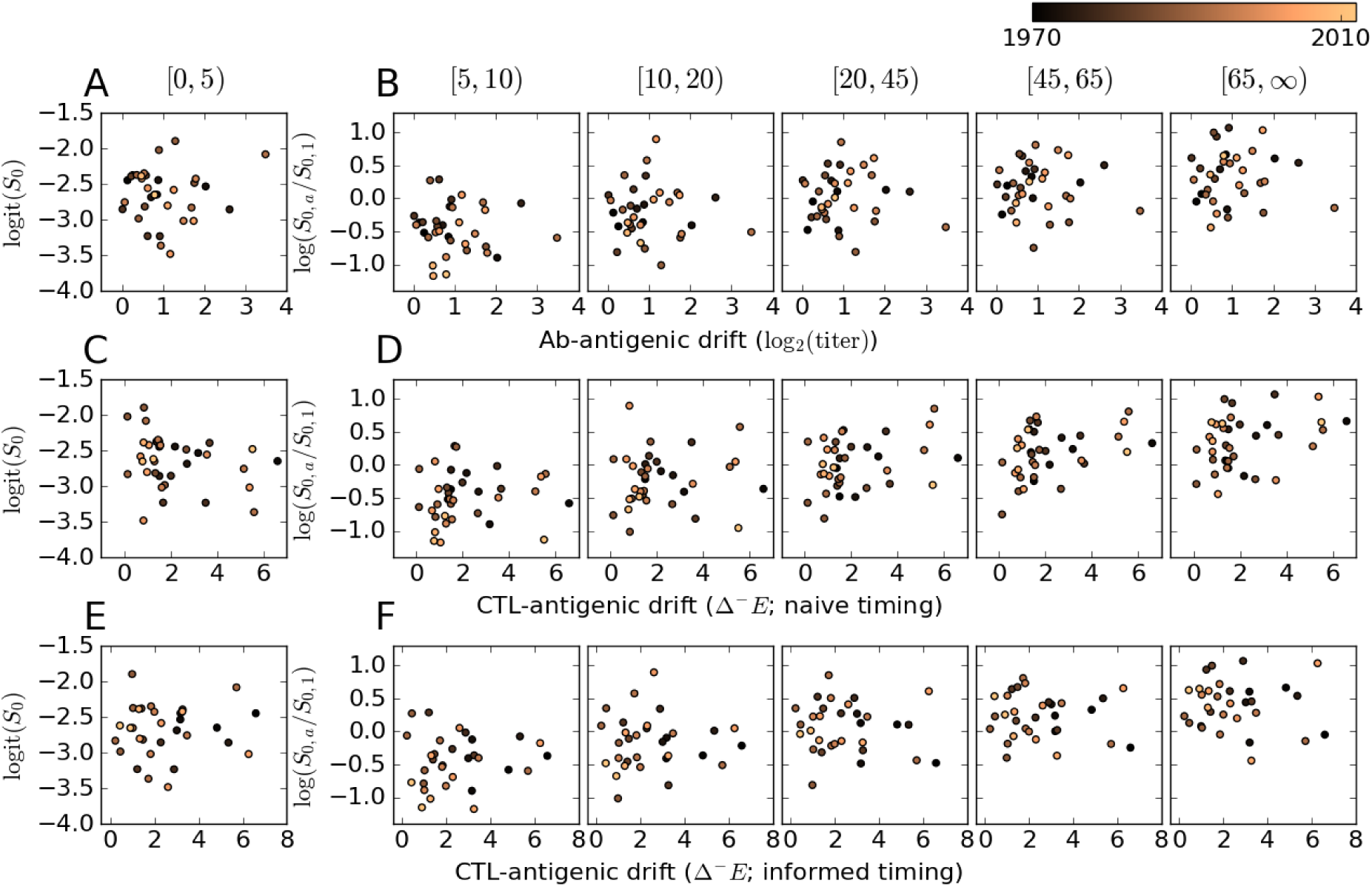
Susceptibility plotted against antigenic drift. Each dot corresponds to an epidemic season (the colors indicate calendar time). For the age class [0, 5), the posterior mean absolute susceptibility is used (A,C,D), and for the other age classes, the posterior mean susceptibility relative to the first age class (B,D,F). We test different antigenic drift values for their predictive power. (A,B) Antibody-antigenic drift, (C,D) CTL-antigenic drift with “naive” timing (i.e. isolates are assigned to a season based on only their calendar year), (E,F) CTL-antigenic drift with “informed” timing (i.e. periods with significant IAV circulation are derived from the fitted model, and missing isolation dates are sampled). The Spearman correlation coefficients are listed in Table 2.

**Table 2:**
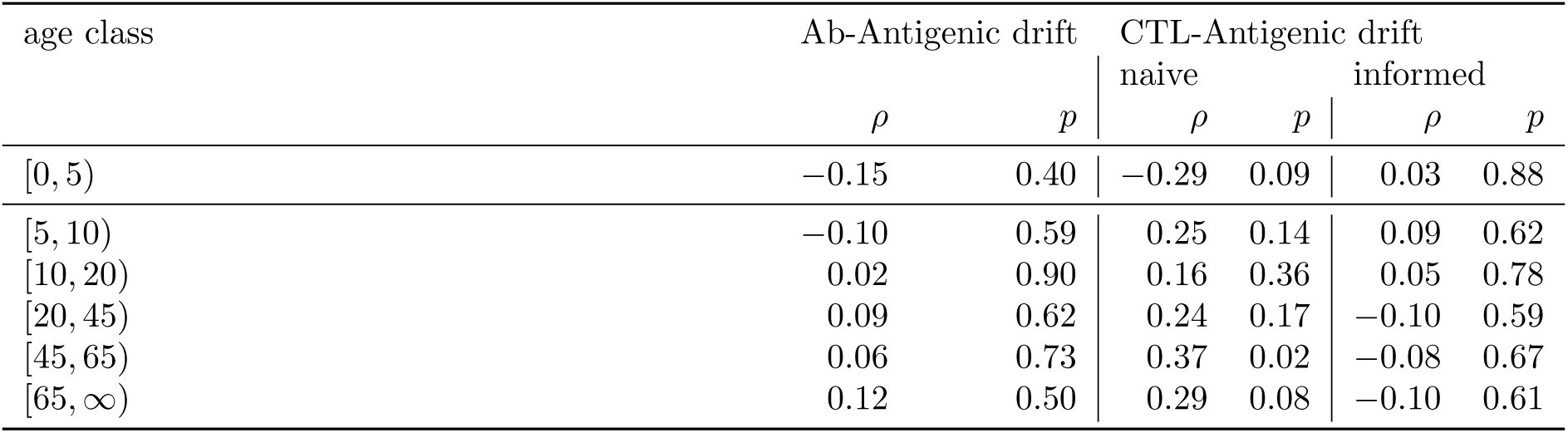
Correlations between antigenic drift and (relative) susceptibility. The correlation is calculated between the posterior mean (relative) susceptibility, and measures of antigenic drift. For the age class [0,5), the absolute susceptibility is used. Listed are the Spearman correlation coefficients (*ρ*), and the corresponding *p-*value. The susceptibility and antigenic drift values are plotted in Figure 6.

We use “epitope loss”, as defined previously (Woolthuis *et al.*, 2016, and see Methods), as a measure for CTL-antigenic drift. Epitope loss measures the number of epitopes that were present in one year, but absent (escaped) in the next. Intriguingly, we do find a significant positive correlation between CTL-antigenic drift and relative susceptibility, but only in the age class [45, 65) (Figure 6B and Table 2). Positive trends can be observed in some of the other age classes, and a negative trend between CTL-antigenic drift and the absolute susceptibility of the youngest age class.

The between-season Ab- and CTL-antigenic drift used in the above analyses has been computed using a number of simplifying assumptions, including the way isolates are assigned to epidemic seasons (i), and what isolates are selected for the analyses (ii). Assumption (i) can be problematic for two reasons. Firstly, the older strains are often only dated at the resolution of calendar years. In the Northern hemisphere, influenza epidemics consistently occur during the winter months, and hence for these strains it is difficult to determine which season they should be assigned to. Secondly, the timing of the epidemic season differs from season to season (Figure 5), in such a way that for some seasons (e.g. 1978/79), the epidemic falls mostly in the first calendar year (1978), while for other seasons (e.g. 1990/91), the epidemic falls in the next calendar year (1991). Assumption (ii) can be problematic when isolates are used that circulate during the summer. Many of them are sampled in the Southern hemisphere, and it is not clear if they can be used as representative strains during the winter epidemic, or for which of the two possible winter epidemics (cf. assumption (i)). Furthermore, sparse sampling of isolates can lead to large uncertainty in antigenic drift.

We resolve both problematic assumptions using a bootstrapping procedure. Using our epidemiological model, we estimate the periods that most likely contains the influenza epidemic (Figure 7A; pink bands). For isolates that have interval censored isolation dates, we sample random dates from a distribution estimated from the known isolation dates (Figure S4). We then sample N = 1000 isolates from the total set of viruses (see Woolthuis *et al.*, 2016) to address assumption (ii), and assign an isolate to a season if it falls into the estimated time interval. Isolates that do not fall into an epidemic season are discarded. This resolves assumption (ii). Using a sampled set of isolates and isolation dates, we re-compute the epitope loss (and gain; see Methods). This procedure is repeated a 1000 times. The result of this more accurately computed epitope loss (and gain) is shown in Figure 7C. Unfortunately, when the timing-aware CTL-antigenic drift replaces the previously used timing-naive CTL-antigenic drift, the previous correlation with relative susceptibility in the age class [45, 65) years is lost (Figure 6F; Table 2).

**Figure 7:**
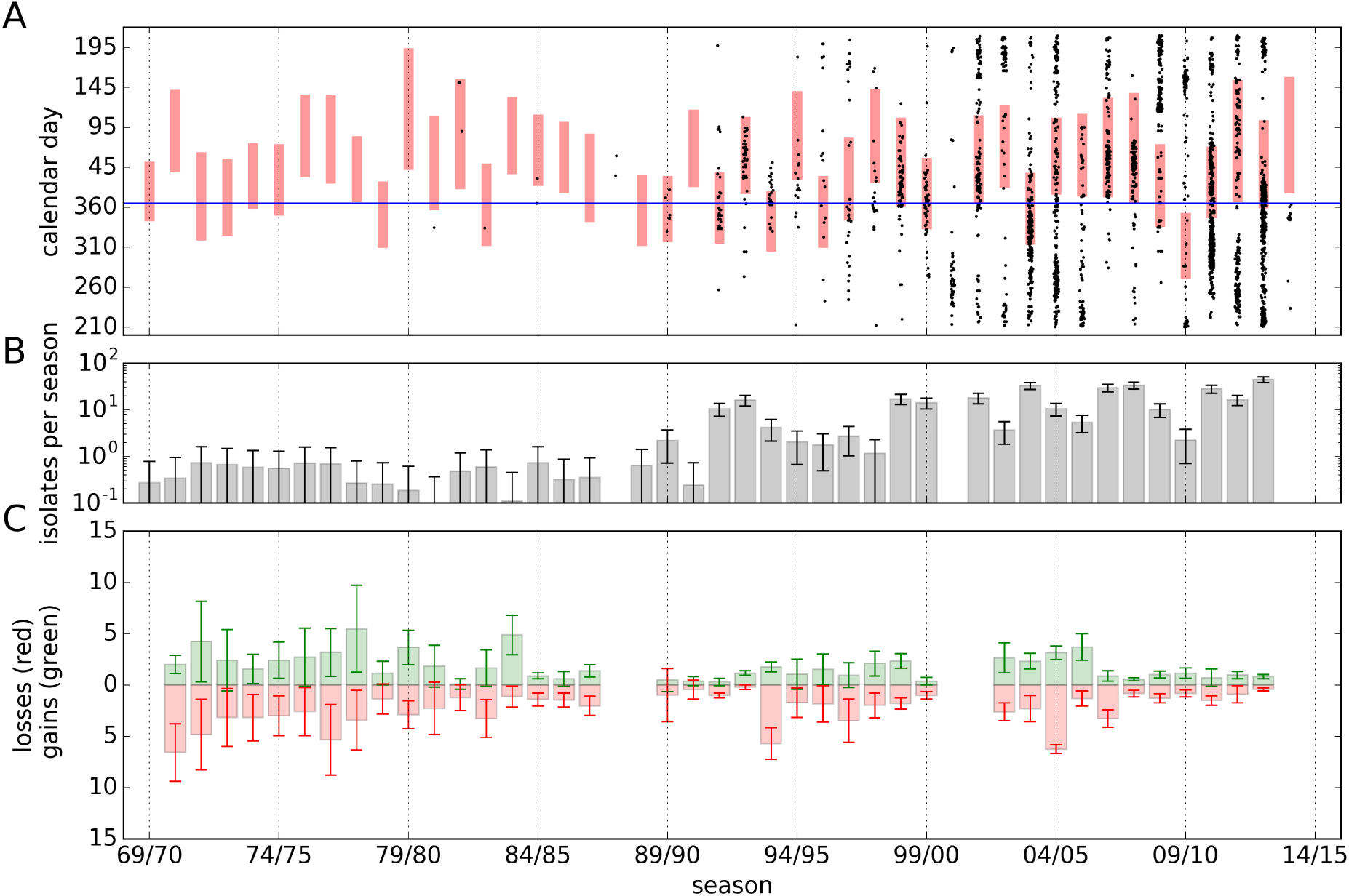
Estimation of CTL-antigenic drift. (A) Time intervals of influenza circulation in the Netherlands, and known sampling dates of the isolates. The red bars indicate the epidemic periods, and the black dots the sampling dates of the isolates. (B) The yearly number of isolates used for the analyses. The whiskers indicate the standard deviation due to the bootstrapping procedure (see Data). (C) The epitope gain and loss, as defined by Equation (1). The whiskers indicate the standard deviation, estimated by bootstrapping.

## Discussion

Using long-term data of ILI reported to GPs, we estimated the yearly age-specific susceptibility of the Dutch population to influenza. Our estimates were obtained using a transmission model in a Bayesian framework, so that all parameters have a clear-cut biological interpretation. We found large variation in susceptibility within age classes, and interesting correlations between age classes that can be explained by expected differences in immunological memory. In contrary to expectation, we were unable to find a relation between Ab- and CTL-antigenic drift in the H3N2 subtype and relative susceptibility.

A large body of work exists where parameters are estimated from ILI (or related) time series. For the Dutch ILI data, this was done by e.g. Xia *et al.* (2005); Te Beest *et al.* (2013); Ratmann *et al.* (2012). Related studies using non-Dutch data were presented by Goeyvaerts *et al.* (2015); Yang *et al.* (2015); Baguelin *et al.* (2013). Most notably, Ratmann *et al.* (2012) use NIVEL’s ILI data in synergy with influenza HA sequence data to test evolutionary hypotheses about the virus. Specifically, using approximate Bayesian computation, they fitted the epochal evolution model of Bedford *et al.* (2012) to the ILI data.

The fact that it seems so difficult to find relations between antigenic drift and susceptibility is curious, and has been noticed before (Van Noort *et al.*, 2012; Chowell *et al.*, 2008). Yet, the belief that variation in attack rates is caused by antigenic drift is widespread. Below, we discuss several reasons for not finding any relation between antigenic drift and susceptibility in detail, but it should be noted that our result, and lack of other epidemiological evidence indicates that the relation between antigenic drift and susceptibility is not as strong and direct as often assumed.

Although we are comparing estimated susceptibility to IAV with disappearance of CTL epitopes, we do not assume that CTL-memory provides neutralizing immunity against IAV infection. More likely, any effect of CTL-memory should be sought in reduced viremia, or enhanced clearance (Gog *et al.*, 2003), and the increased likelihood of asymptomatic infection (Hayward *et al.*, 2015). If asymptomatic infection coincides with decreased transmission, asymptomatically infected individuals can be counted as non-susceptible, since a model with an asymptomatic compartment is equivalent to the usual SIR model (Van Noort *et al.*, 2012, and Supplementary material).

We compared relative susceptibility with a simple measure of antigenic drift. In reality each individual will have its particular immunity against future influenza strains, due to the many influenza infection histories that are possible (Fonville *et al.*, 2014), and therefore each individual experiences the antigenic drift or shift differently (Cobey and Pascual, 2011; Cobey and Hensley, 2017). As an extreme example, HA-imprinting and the replacement of H1 with H2 in 1957 and then H3 in 1968 is likely the reason why birth year explains susceptibility to either severe H7N1, or severe H5N1 avian influenza infections (Gostic *et al.*, 2016). Because we consider T-cell responses, HLA polymorphism complicates the matter even further; each HLA haplotype results in a different set of possible epitopes. Possibly, an agent-based model can be used to generate individual infection histories, giving a more complete measure of the effects of CTL-antigenic drift, that can then be compared with susceptibilities measured from ILI data. However, such an approach is not straightforward. For instance, as mentioned by Ratmann *et al.* (2012), estimation of the effect of immune histories on susceptibility is sensitive to small uncertainties in the historical sequences and ILI data. Our previous considerations about epitope loss indicate that these uncertainties may not even be that small.

Another source of uncertainty is the fact that ILI is not a very specific predictor for influenza. The positive predictive value (PPV) of ILI has been estimated at ≈ 50% for influenza during the peak ILI weeks of the seasons 2003/04 to 2011/12. We attempt to correct for this relatively low PPV by using a phenomenological model for the background ILI. When instead the background ILI is assumed to occur at a constant rate throughout the year (the “constant background ILI” scenario; Table 1), the estimates of *ℛ*_eff_ tend to stay away from the epidemic threshold (min *ℛ*_eff_ = 1.11), since background ILI wrongfully has to be explained by influenza. However, we expect some model misspecification by ignoring the fact that background ILI, akin to influenza, is caused by infectious pathogens.

Our assumption about the mechanism underlying the trend in the reporting rate is consistent with other studies (Dijkstra *et al.*, 2009), but not based on hard evidence. Although our estimates for more recent years are consistent with an independent data source (GIS), the high reporting rate during the seventies could be an artifact of our model. Van Noort *et al.* (2012) have hypothesized that not infectiousness, but reporting rate is influenced by weather, and use this to explain variation in reported incidence. However, such a hypothesis is not sufficient to explain the strong decrease in reported ILI during the last five decades. In a later study (Van Noort *et al.*, 2015), differences in reporting rate between countries are attributed to cultural differences. The differences in reporting rate between countries are of the same order of magnitude as the difference within the Netherlands between 1970 and 2014. Another possibility is formulated by Morens *et al.* (2009), who hypothesize that for evolutionary reasons, the virulence of influenza could be decreasing, while infectiousness is retained; a host that is not put to bed will spread a disease more efficiently. Although intriguing, this idea does not fit well with our model, since not only influenza reporting decreases, but also reporting of background ILI. Finally, the fitness of seasonal influenza could be slowly deteriorating, possibly due to continuous antigenic drift needed for immune evasion, as argued by Fleming and Elliot (2008) based on an ILI time series from England and Wales. However, following the same reasoning as above, most ILI-causing pathogens should then be subject to fitness erosion. This includes (in hindsight) the pandemic H1N1 virus from 2009/10, since the incidence during this season is not strikingly different from its surrounding years (Figure 1).

Some of the decreasing ILI in the age class [65, ∞) could be the result of vaccination policy (McDonald *et al.*, 2016) and uptake (Spruijt *et al*, 2016). As of the year 1997, vaccination against influenza for individuals in this age class is free of charge in the Netherlands. Despite the fact that we use a trended *q* to account for the decreasing ILI, one might still expect a difference in susceptibility before and after 1997 for the elderly. However, we cannot find evidence of an effect of vaccination by making this comparison (*p* = 0.47, *t*-test based on posterior means). Analogous to antigenic drift and susceptibility, the difficulty with finding associations between vaccine uptake and ILI incidence may be due to complex infection histories itself (Spruijt *et al*, 2016),

Recapitulating, our analyses have shown that, on the one hand, there are systematic and substantial differences between age classes in their susceptibility to IAV infection, with 5 to 20 year-olds being least susceptible, and infants and young children ([0, 5) years) and older adults (65 years and older) being most susceptible. On the other hand, there is also substantial variation in susceptibility between years, thereby precluding attempts to quantitatively predict age-specific susceptibility with meaningful precision. Although it is generally believed that these patterns of differences in susceptibility between years and between age-groups are molded by viral evolution and pre-existing immunity, we were unable to find evidence for this hypothesis in our data. This was true both for humoral immunity mediated by antibodies directed against the hemagglutinin protein and also for cellular immunity mediated be cytotoxic T cell responses. Hence, we are led to the perhaps somewhat sobering viewpoint that the complex interplay of viral evolution and pre-existing immunity is highly sensitive to details of the infection histories in narrow age strata (Ratmann *et al.*, 2012; Gostic *et al.*, 2016), and may be fundamentally unpredictable.

## Methods

### Data

We made use of a number of data sources. For self-consistence, we here briefly discuss these sources, and our particular usage of the data.

### NIVEL

The catchment population covers approximately 1% of the Dutch population, equally distributed over different regions in the Netherlands (Donker, 2016; Dijkstra *et al.*, 2009). Using the ISO week date system, we translated years and week numbers into the number of days since January 1, 1970. This solves the issue of ‘leap weeks’, which should be treated with care, since they tend to coincide with the epidemic peaks.

The data is stratified into 6 age classes: [0, 5), [5, 10), [10, 20), [20, 45), [45, 65) and [65, ∞) years, usually indexed by *a* = 1,…, 6, respectively. As of 1986, the resolution of the age stratification has been increased. Our age classes are chosen to be compatible with early and more recent ILI data stratification, and to fit well with contact-intensive clusters (not shown) that can be observed in the contact matrix (see below). Some of the ILI data is missing, but has been partially recovered (see Missing data).

Dutch law allows the use of electronic health records for research purposes under certain conditions. According to this legislation, neither obtaining informed consent from patients nor approval by a medical ethics committee is obligatory for this type of observational studies containing no directly identifiable data (Dutch Civil Law, Article 7:458). This study has been approved by the applicable governance bodies of NIVEL Primary Care Database under nr NZR00316.056.

### POLYMOD

The POLYMOD study is a prospective survey of social contact patterns in multiple countries (Mossong *et al.*, 2008; Wallinga *et al.*, 2006). The key assumption for epidemiological studies of respiratory-spread infectious diseases is that conversational contact can be used as a proxy for infectious contact (the social contact hypothesis, Wallinga *et al.*, 2006). Our contact matrix is based on only the Dutch data, and estimated using a method developed by Van de Kassteele *et al.* (2017). In the study, *participants* (see Figure S2) were asked to keep a diary on the individuals with whom they had *contact* during one day. The average numbers of contacts applied to our age classes are given in Figure S2.

### GIS

The Great Influenza Survey is the Dutch branch of the European Influenzanet project (Van Noort *et a*l., 2015), a monitoring system for **ILI,** using voluntary cohorts approached via the Internet. In the on-line questionnaire, the participants are asked, among other things, if they had influenza-like symptoms such as fever and coughing, and if they consulted a GP while they had these symptoms.

Most of the GIS data used in this paper is published by Friesema *et al.* (2009). Unfortunately, the age stratification in this report has a slight mismatch with our choice of age classes, and in order to compute confidence intervals, the population sizes of self-diagnosed ILI patients, and the sub-population seeking a GP consult are needed. I. Friesema kindly provided the original data, which enabled us to compute the population sizes for our age stratification.

### Epitopes

The full procedure for collecting epitopes is described in (Woolthuis *et al.*, 2016). In short, we collected all complete, human IAV proteomes without ambiguous amino-acids from the GISAID EpiFlu database (www.gisaid.org). All known CTL epitopes were downloaded from IEDB (www.iedb.org). Epitope sequences that contained shorter epitopes were filtered out. For each epitope *e* and each calendar year *y*, the fraction of IAVs that contained the epitope is denoted by ϕ_y,e_. Epitope gain 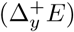 and loss 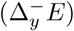 were defined as

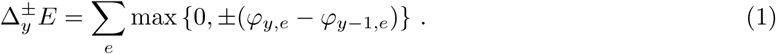

In order to estimate the uncertainty of the gain and loss, we applied the following bootstrap procedure. First, we sample (with replacement) a 1000 strains from the total number of 3321 isolates. Using the sample, we compute gains and losses (Equation 1) for every year *y*, whenever the sample contains isolates from both years in the pair *(y –* 1, *y).* We repeat the sampling and computation *n =* 1000 times. Notice that we produce less bootstrap samples for a year *y*, whenever few strains were isolated in the year *y* or the previous year *y –* 1.

In order to assign isolates to seasons in an informed manner, we first have to estimate when the epidemics took place. We assign a a week to the epidemic, when the weekly incidence of the total catchment population exceeds 10 cases per 100 000 individuals. Hence, when 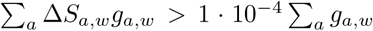 (using notation introduced below). We then define *t*_*start*_ as the first day of the first epidemic week and *t*_*end*_ as the last day of the last epidemic week. Notice that *t*_start_ and tend need not be defined; when R*_eff_* is close to 1, the incidence may never reach the threshold. This leads to epidemic periods that are on average 12.3 weeks long (IQR: [11.0,14.0] weeks). On purpose, our the epidemic threshold is 5 times smaller than the guideline used by NIVEL (51 cases per 100 000), resulting in slightly longer epidemics. Isolates are then assigned to season s, if they were sampled within the epidemic period [*t_start_,t_end_*] of season *s.* Isolates that do not fall into any epidemic period are ignored. Missing isolation dates are sampled from a distribution inferred from the known dates (Figure S4). This distribution is estimated by fitting a Circular-Gaussian kernel with a bandwidth of 30 days to the set of known calendar days. The bootstrapping procedure is similar as in the naive case. A 1000 strains are sampled (with replacement), then, sampled isolates with missing dates are randomly generated, and finally we assign seasons to the isolates. Epitope gain and loss is computed as usual.

### Dynamical model

We model the influenza epidemics using a system of ordinary differential equations (ODEs). The population is partitioned into classes depending on disease status: susceptible (S), infectious (I) and recovered (R). We use an age stratified ‘SIIR’ model, with fixed contact rates *C* (Figure S2). The model is given by the following equations:

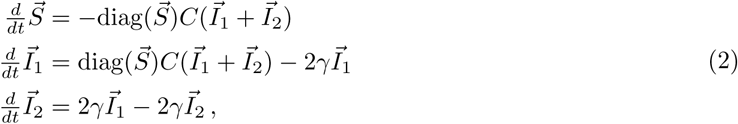

where diag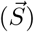 is the diagonal matrix with the vector 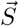 on the diagonal. The two infectious phases are used to better model (non exponential) distribution of the length of the infectious period. The initial condition at time *t = t*_0_, is given by a small perturbation (of size ε, with 0 < ε = 10–^6^ ≪ 1) of the disease-free steady state: 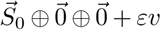, where 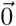 is the zero vector in ℝ^6^, and ⊕ denotes concatenation of vectors. Here, *v* is the dominant eigenvector of the Jacobian of system (2) at the state *S*_0_ © 0 © 0. The vector *v* has positive I-coordinates and satisfies ||*v||_2_ =* 1. Using the S-coordinates of the solution for (2), we compute the weekly incidence 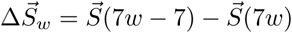.

The likelihood of observing *f*_*aw*_ ILI cases in age class *a* during week *w*, is assumed to be Poisson distributed:

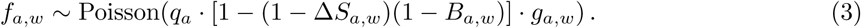

The Poisson distribution is taken because it approximates the Binomial distribution, but can be computed more efficiently, and has convenient additive properties (see below). The expectation is the product of the catchment population size *g*_*a,w*_, the reporting probability *q*_*a*_, and the probability of contracting either influenza *(∆S*_*a*,w_), or another ILI-causing pathogen *(B*_*a*,w_).

The incidence of background ILI *(B_aw_)* is modeled using the sine function, in order to capture seasonal effects (cf. Van Noort *et al.*, 2012). We set *B_a,w_ = B_a_(7 w)*, where

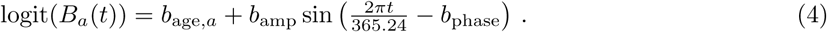

The age effect *b*_*age*_, the phase *b*_phase_ ∈ [0, 2π), and the amplitude *b*_amp_ ∈ [0, ∞) of the background ILI have to be estimated.

The reporting probability *q*_*a*_ is allowed to follow a trend, by defining

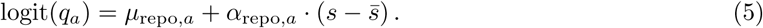

The logit function on (0,1) is defined by logit(*x*) = log(*x*/(1 – *x)).* The seasons (*s*) are centralized (by subtracting the middle season *s =* 22 from s) to reduce the correlation between *µ*_*repo*_ and *α*_*repo*_

### Estimation of the parameters

The parameters of the model are estimated using Markov chain Monte Carlo (MCMC). Since ODEs have to be integrated in order to compute the likelihood, we implemented a (Metropolis within) Gibbs sampler in C++. The code, together with mock data and a Python script for interpreting the output, has been made publicly available (Van Dorp, 2016). The costly likelihood computation can be accelerated by updating the seasonal sub-models in parallel, and using the fact that updating of many of the parameters does not require re-integration of the ODEs. The length of the chains is 2 x 10^5^, and only the second half is used in the analysis, after applying a 1 : 100 thinning. Convergence of the chain was assessed visually, by inspection of the trace plots.

The susceptibility parameters *S*_0,_*_a_* are given Logit-Normal prior distributions, dependent on season and age class a, with mean 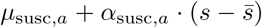 (where *s* denotes the season, and 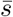 the middle season) and variance 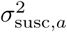, which are given non-informative hyper-priors *µ*_*susc*_ ~ Normal(0,100), *α*_*susc*_ ~ Normal(0,100) and 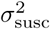 ~ Half-Normal(100). In this way, susceptibility during seasons, or for age classes with little information, are informed by the other seasons and age classes. Similarly, the (relative) onsets of the epidemics *t*_0_ are given a Normal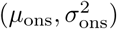 prior, with hyper-priors Normal(100,1000) ~ *µ*_ons_ and Half-Normal(1000) ~ 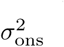. The average background ILI b_mean,a_ is given a Normal(0,100) prior, and the parameters governing the seasonality of the background ILI rate, *b*_*amp*_ and *b*_phase_, are given a Half-Normal(100) and a Circular-Uniform(0, *2*π) prior, respectively. The offset µ_repo_ and slope α_repo_ of the logit-reporting rate both have a Normal(0,100) prior distribution.

The proposal distribution for each parameter is given by a symmetric mixture of normal distributions (Yang and Rodriguez, 2013), with mean equal to the present state of the parameter. The proposal variance is tuned during the burn-in phase to achieve an acceptance rate of about 0.44 (Rosenthal, 2011). For parameters whose domains have boundaries, the proposal is reflected in these boundaries to make sure that the Markov chain converges to the posterior distribution (due to the ‘detailed balance’ condition). Likewise, a wrapped version of the proposal is used for circular domains.

### Missing data

Some ILI data is missing (see the gray blocks in Figure 1), but part of this data could be retrieved from (Xia *et al*., 2005). This recovered data is, however, not age-stratified, and also the catchment population sizes *g* are absent; instead the fraction of ILI incidence is reported. In order to compute the likelihood of observing a fraction of ILI incidence, given our model, we took the following steps. First, the time series of the present catchment population sizes shows a highly predictable pattern (Figure S1), and hence we took the simple approach of filling in the missing population sizes by taking a weighted average of the values in the 4 surrounding years:

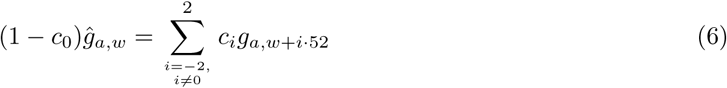

with *c_±2_ =* 0.061, *c*_±1_ = 0.245, and *c*_0_ = 0.388 (a Gaussian kernel). Second, suppose that *µ*_*a*_ is the expected ILI incidence (see Equation 3), and π is the reported fraction of ILI incidence, and *f*_*a*_ is the true ILI incidence, then 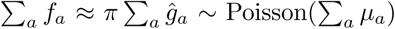, because of the additive properties of the Poisson distribution. The Gaussian kernel model does not correspond to the data that well between the years 2004 and 2006, but this poses no problem, since the data is age-stratified around that time.

### Model selection

We compare the different models using the WAIC (Watanabe, 2010; Gelman *et al*., 2014), and the WBIC (Watanabe, 2013). Models with a lower WAIC or WBIC are able to describe the data better. The WAIC, is an estimate of the out-of-sample prediction error. Ideally, one would use Bayes factors to compare different models, but computing Bayes factors is highly impractical in our case. The WBIC is acts as an approximation for the marginal likelihood (and hence, the difference in the WBIC for the Bayes factor). For non-singular models, one could replace WAIC by AIC, and WBIC by BIC. However, as our model is singular due to the use of hyper-parameters, we need to use WAIC and WBIC.

The WAIC equals 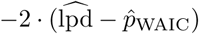, where 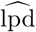 is the log point-wise predictive density, and 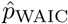 is a penalty term based on the effective number of parameters (Gelman *et al.*, 2014). Unlike the WAIC, the WBIC can not be computed from the output of the Gibbs-sampler. Instead, sampling has to occur at a different “temperature” determined by the number of observations (*n* = 13333). More precisely, 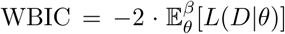, where *L*(*D*|*θ*) equals the log-likelihood of the data *D* given the parameters *θ*, and 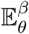 denotes the expectation with respect to the distribution function proportional to 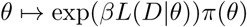, with π denoting the prior distribution of *θ*, and *β =* 1/log(*n*) ≈ 0.11.

## Acknowledgements

We gratefully acknowledge Ingrid Friesema for helping us with the GIS data and Jan van de Kassteele for computing a customized contact matrix. We thank Dennis te Beest and Johannes Textor for technical support, and Jantien Backer for comments on an earlier version ofthe manuscript. Finally, we are grateful to Mariette Hooiveld (NIVEL Primary Care Database, Sentinel Practices, Utrecht, The Netherlands) for her assistance with the ILI data and her comments on an earlier version of this manuscript.

This research was funded by NWO (www.nwo.nl, grant numbers 645.000.002 and 823.02.014) and the National Institute for Public Health and the Environment (www.rivm.nl).

## Supplementary material

### Models with asymptomatic infection

CD8^+^ T-cell memory might not result in reduced susceptibility, but does lead to less severe disease, and in the best case asymptomatic infection. Therefore, a model that includes an asymptomatic compartment would better fit with our assumptions about the epidemiology of influenza. However, as pointed out before for a simple case (Van Noort *et al*, 2012), such a model is equivalent to the usual SIR model, and the additional parameters are not identifiable.

Since we use a slightly more intricate model than the one explored by Van Noort *et al.* (2012), we here show that also our model is equivalent with a version that includes asymptomatic compartments. This version of the model is given by the following ODEs:

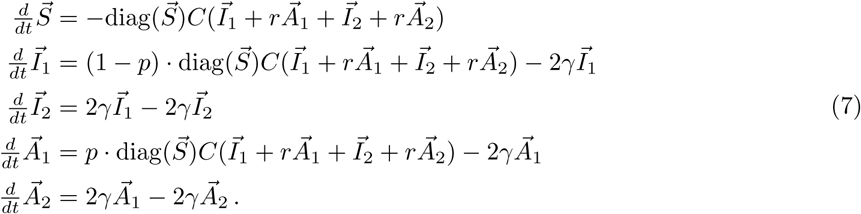

This model has two additional parameters: the reduction of infectiousness of the ‘asymptomatic’ individuals in the A compartment (*r*), and the fraction of susceptible individuals that move to the A compartment upon infection (*p*). We have added 2 compartments A_1_ and A_2_ (of size *A*_*1*_ and *A*_*2*_, resp.) such that the asymptomatic phase has the same duration as the infectious phase.

When *r* = 0, individuals in compartment A are not infectious (possibly due to largely reduced viral shedding). In this special case, we can simply ignore the equations for 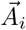. We will not be able to estimate the parameter *p.* Instead, we can write 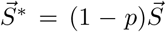, and can write the system of ODEs in terms of 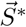, and 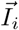. The variable 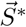 can then be interpreted as the fraction of individuals that are susceptible for *symptomatic* infection. For general 0 ≤ r ≤ 1, we can write 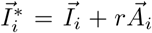, and define 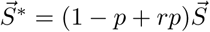, so that

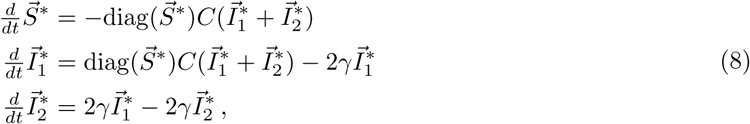

and these equations are identical to system (2). It is not unreasonable to assume that individuals in the A compartment have a reduced probability of consulting their GP compared to individuals in the I compartment. Let us denote the reporting probabilities *q*_*a*_ and *q*_*I*_, respectively. The rate of reported ILI is therefore 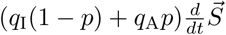. If we further assume that the probability of reporting is proportional tothe reduction in infectiousness, we get that 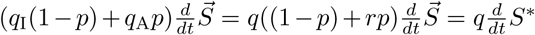 Hence, under these assumptions, the baseline model (2) is equivalent to a model with asymptomatic infection, but with *S* interpreted as susceptibility weighted by the likelihood of seeking health-care.

### Models with frailty

In the case of SIR-type models, the term “frailty” refers to heterogeneity in susceptibility. The two extremes are zero-one susceptibility (a person is either fully susceptible to infection or not at all) and uniform susceptibility (everyone has the same (reduced) susceptibility).

An example of a model that incorporates both of these susceptibility notions is given by the following equations:

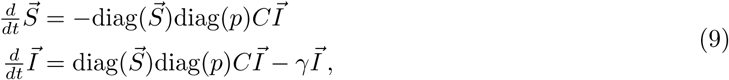

with initial condition 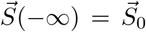. Here, the parameter *p* denotes the reduced susceptibility of the different age classes. Let 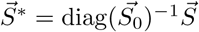, and 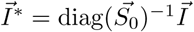, then

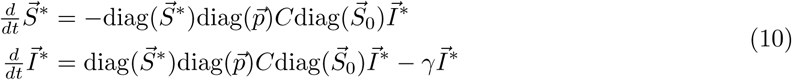

The force of infection acting on age class a, is given by

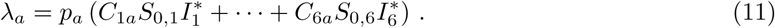

Above, we interpret the parameters *S*_*0,a*_ as the fraction of susceptible individuals (by using 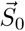 for the initial condition). Equation (11) shows that *S*_*0,a*_ can equally well be interpreted as reduction in infectiousness. In the case of immunity due to neutralizing antibodies, the best interpretation of *S*_*0,a*_ would be the fraction of susceptible individuals. For cellular immunity, *S*_*0,a*_ can probably be best interpreted as reduced infectiousness of individuals in age class *a.* Hence, in general, *S*_*0,a*_ should be interpreted as a combination of both.

**Supplementary Table 1:** The posterior modes (MAP estimates) and the 95% CrIs of the susceptibilities (*S*_0_), the effective reproduction numbers (*ℛ*_eff_), and the onset of the epidemics (*t*_0_), are listed in the supplementary file `SupplTable.tsv` as tab-separated values.

---

**Figure S1:**
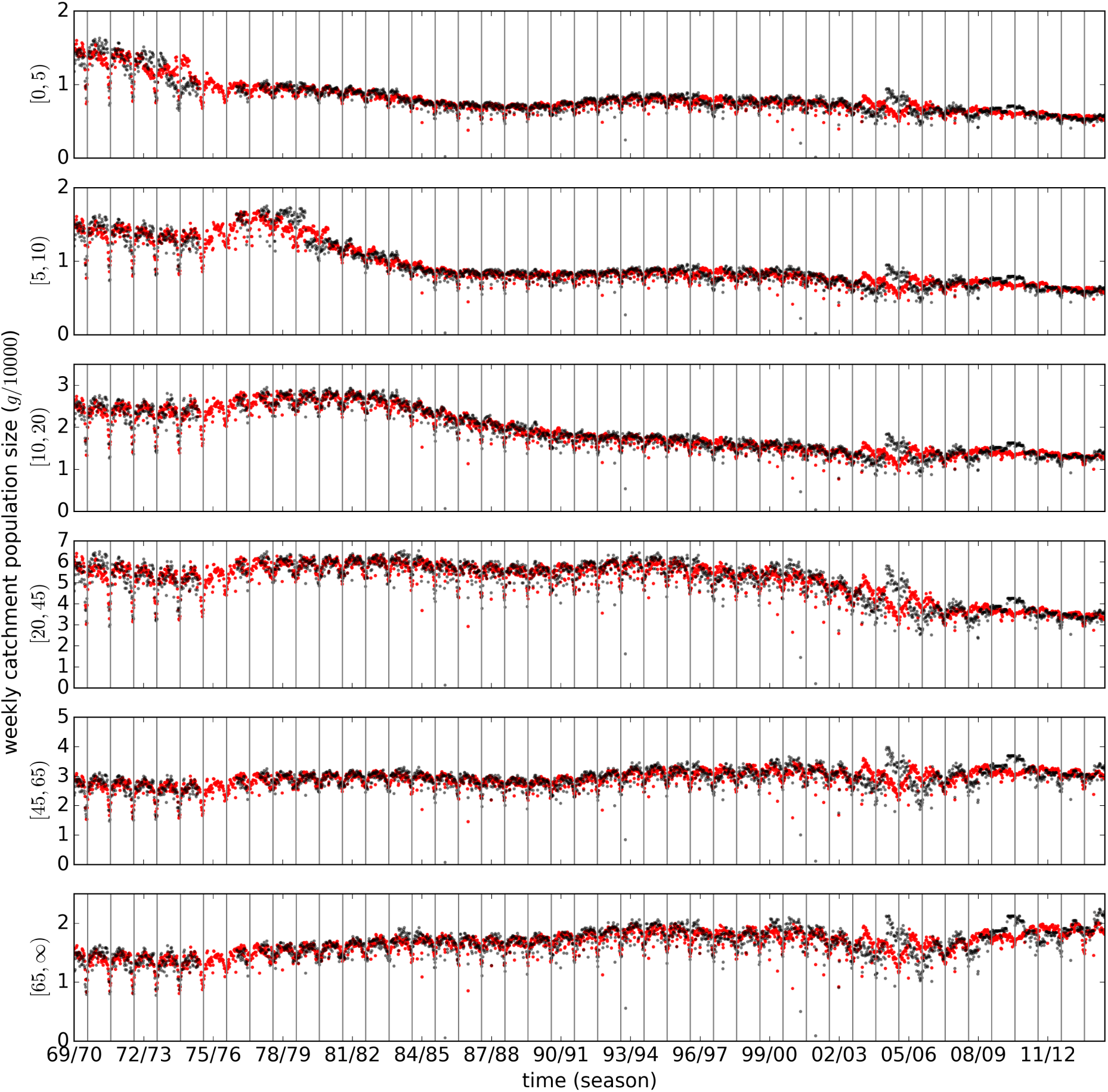
Catchment population size. The actual population sizes g are shown in black. The estimates (gˆ; Equation 6) based on the 4 surrounding years are shown in red. Only when the population size is not known, an estimate is used instead. The vertical lines show the beginning of the seasons (week 30).

**Figure S2:**
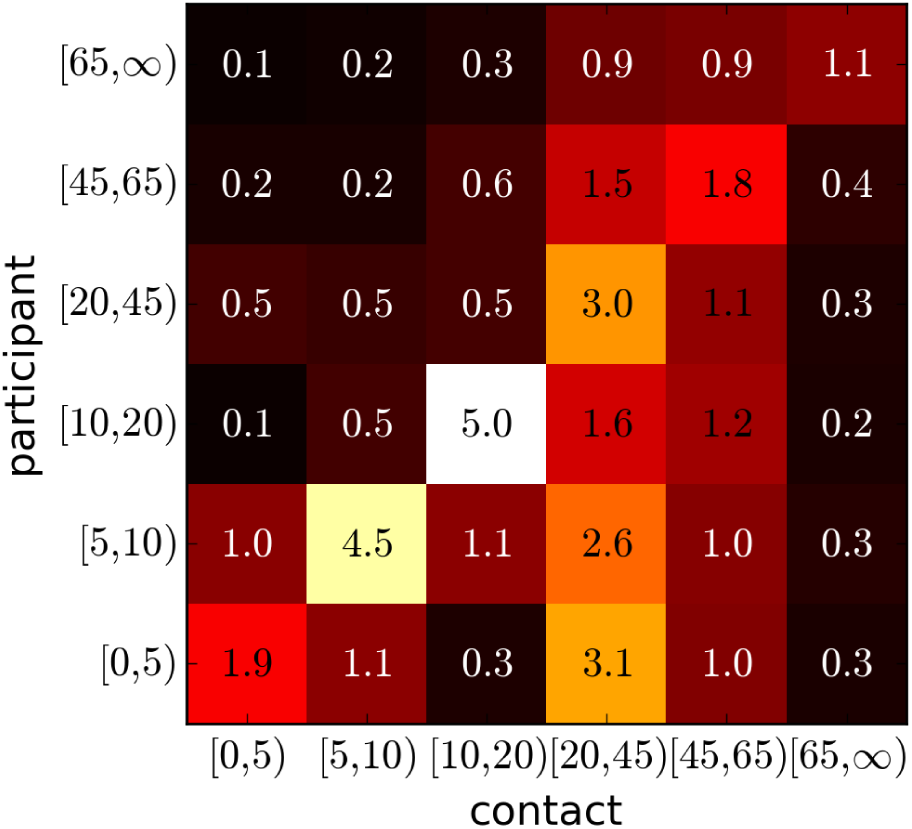
Contact matrix (C). The values represent the average number of contacts per day. The dominant eigenvalue of C equals 7.1.

**Figure S3:**
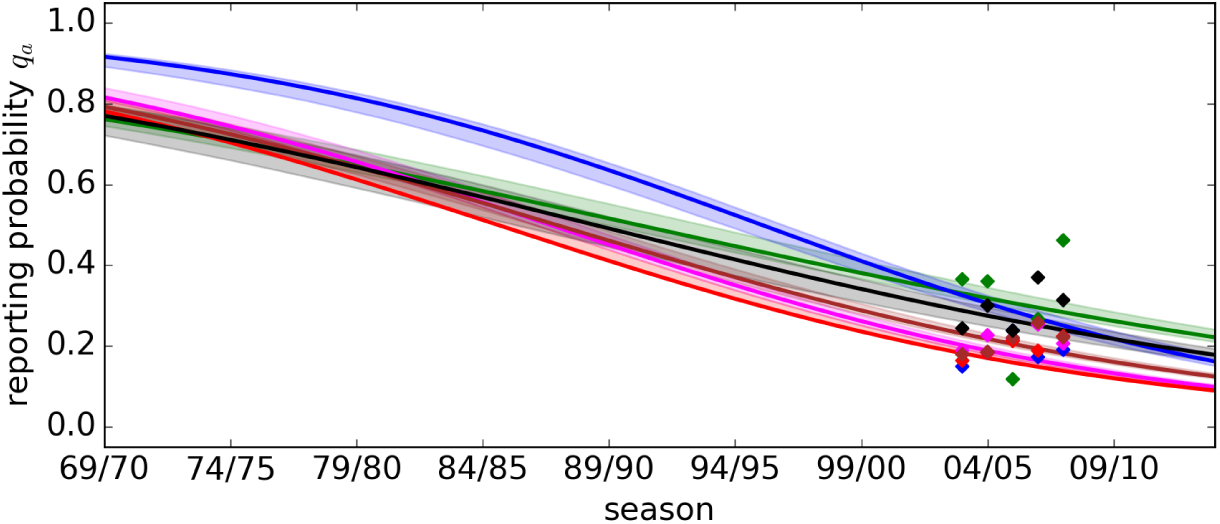
The reporting probability decreases with time. The reporting probabilities are given as a function of time (lines for the MAP estimate, and bands for the 95% CrI). The color coding for age class is identical to Figure 3. Notice that during a season the reporting probability is kept constant. The diamonds give the estimates based on the GIS data (cf. Figure 2).

**Figure S4:**
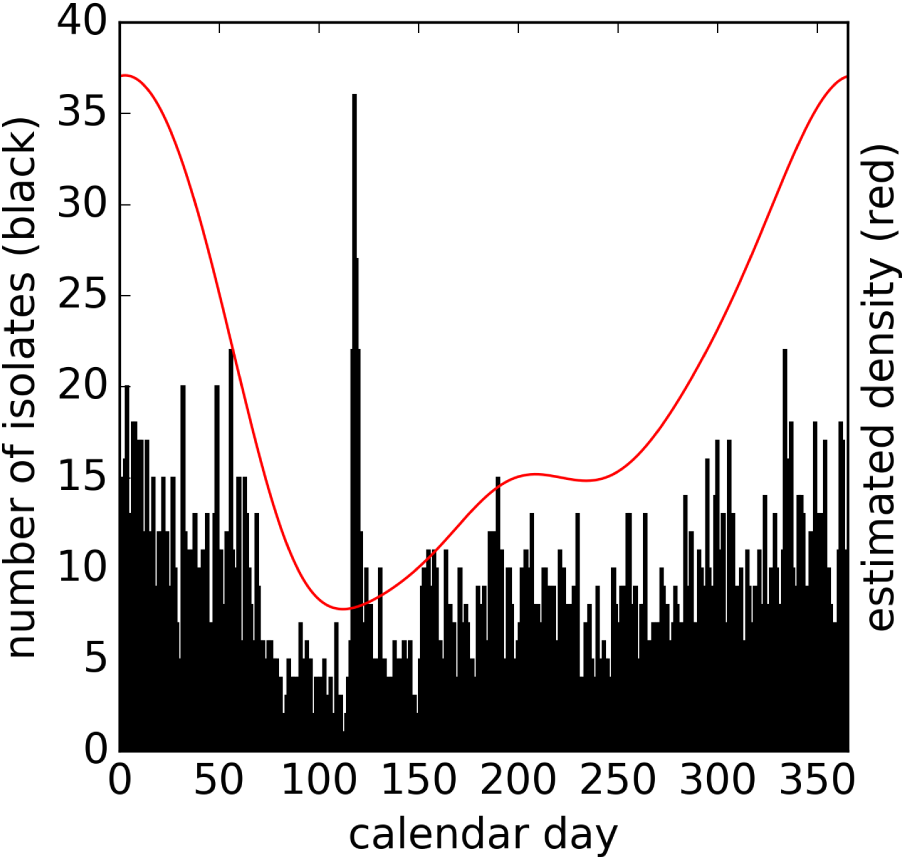
Distribution of isolate sampling dates. The distribution of known isolate sampling calendar days is shown as a black histogram. The red line represents the smoothened density function.

